# Novelty Processing Depends on Medial Temporal Lobe Structures

**DOI:** 10.1101/2020.11.10.374538

**Authors:** J. Schomaker, M.M.E. Grouls, E. Rau, M. Hendriks, A. Colon, M. Meeter

## Abstract

**Objectives:** The goal of the present study was to identify the role of the medial temporal lobe (MTL) in the detection and later processing of novelty stimuli.

**Methods:** Twenty-one epilepsy patients with unilateral MTL resection (10 left-sided; 11 right-sided) performed an adapted visual novelty oddball task. In this task two streams of stimuli were presented on the left and right of fixation while the patients’ electroencephalogram was measured. Patients responded to infrequent target stimuli, while ignoring frequent standard, and infrequent novel stimuli that could appear either contra- or ipsilateral to the resected side.

**Results:** Novelty detection, as indexed by the N2 ERP component elicited by novels, was not affected by the MTL resections. Later processing of novels, however, as indexed by the novelty P3 ERP component, was reduced for novels presented contra-versus ipsilateral to the resected side. Target processing, as indexed by the P3b, was unaffected.

**Conclusions:** The current results suggest that MTL structures play a role in novelty processing, but that the novelty signal may originate from a distinct neural source.

## Introduction

The human brain can very quickly distinguish between new and familiar stimuli. Novelty signals have been reported as early as 85 milliseconds after stimulus presentation, when recognition of old or new information is rewarded (Bunzeck, Doeller, Fuentemilla, Dolan, & Duzel, 2009). Even without added motivational value, novelty signals have been identified within 150 ms post stimulus (Düzel, Habib, Guderian, & Heinze, 2004).

A novelty signal is believed to originate from the hippocampus, a MTL structure (Knight, 1996; Kumaran & Maguire, 2009; Lisman & Grace, 2005; Ranganath & Rainer, 2003). The involvement of the MTL in creating new memories has been known for more than a century (Bechterew, 1900), and explicit recollection depends on the MTL (Squire, 1992; Squire, Stark, & Clark 2004). Patients with bilateral damage to the MTL cannot convert new experiences into memories that can be consciously recollected later on (Penfield & Milner, 1958; Milner, 1972; SCOVILLE & MILNER, 1957). As learning and identifying new stimuli go hand in hand, the role of the MTL in detecting and processing new events has been assumed for a long time. Indeed, neuroimaging studies (Bunzeck & Düzel, 2006; Kaplan, Horner, Bandettini, Doeller, & Burgess, 2014; Murty, Ballard, MacDuffie, Krebs, & Adcock, 2013; Wittmann, Bunzeck, Dolan, & Düzel, 2007) and electrophysiological recordings (Fried, MacDonald, & Wilson, 1997; Grunwald, Lehnertz, Heinze, Helmstaedter, & Elger, 1998) in humans have confirmed that the MTL, and specifically the hippocampus, is involved in processing of new and unexpected information. Results from an fMRI study showed that the anterior hippocampus responds to stimuli that are either perceptual, semantic, or emotional oddballs (Strange & Dolan, 2001). The authors argued that the anterior hippocampus detects a general mismatch between expectations and outcome, rather than novelty per se. Converging evidence thus suggests that the MTL, and specifically the hippocampus, is required for the detection of unexpected, deviant, and/or novel information (Aito, Abbate, Marcucci, & Cominelli, 2007; Kumaran & Maguire, 2009).

Traditionally, physiological novelty responses have been studied using a three-stimulus novelty oddball paradigm and the event-related potential (ERP) technique. In the visual version of this task infrequent task-irrelevant novel images are presented in a sequence of frequent standard, and infrequent target images. The novels elicit a frontal negative ERP component between 200 and 300 milliseconds, which has been referred to as the anterior N2, novelty N2, and sometimes N2b (Folstein & Van Petten, 2008), believed to reflect novelty detection, or the stimulus-driven perceptual part of processing (Ferrari, Bradley, Codispoti, & Lang, 2010; Schomaker & Meeter, 2014; Schomaker, Roos, & Meeter, 2014). This negative component for novels is typically followed by a positive-going ERP component peaking slightly later between 300 and 450 ms over frontocentral regions, which has been called the novelty P3 or P3a (Friedman, Cycowicz, & Gaeta, 2001; Schomaker et al., 2014). The novelty P3 and P3a were originally believed to reflect different cognitive processes; the novelty P3 being used as an index of the orienting response to novel stimuli, while the P3a was associated with processing attended and unattended deviant stimuli (Squires, Squires, & Hillyard, 1975). Some have failed to distinguish the two (Hay et al., 1995), while others still argue they reflect different processes (Barry, Steiner, & De Blasio, 2016). Here, we will use the term novelty P3, to refer to the novel-elicited positive component peaking after the N2. The novelty P3 can be distinguished from the P3b (or P300) component elicited by target-relevant stimuli and peaking slightly later over posterior regions.

The ERP technique has been used for over 40 years to investigate the brain’s response to novelty. While its spatial resolution is poor, it still has a higher temporal resolution than newer neuroimaging techniques, having the advantage that different aspects of stimulus processing can be disentangled. For example, the earlier novelty N2 component has been associated with detection of novelty (Schomaker & Meeter, 2014; Schomaker et al., 2014), while the later novelty P3 component has been associated with the conscious evaluation of a novel stimulus (Friedman et al., 2001), and detection of deviance (Barkaszi, Czigler, & Balázs, 2013; Schomaker et al., 2014). Some studies have suggested that the P3 signal originates from the hippocampus. Using intracranial recordings, Brankack, Seidenbecher & Müller-Gärtner (1996) observed P3-like components in the neocortex but with peaks in the CA1 region of the hippocampus in awake rats to novel stimuli during an auditory discrimination task, akin to the auditory oddball task used in humans. Hippocampal evoked responses to novel stimuli could already be observed within the first 100 milliseconds post-stimulus. They argued that the P3 component originates from initial bursts of activity within the hippocampus, suggesting that a novelty signal may indeed be generated in the hippocampus.

Consistent with this, the study of Rutishauser, Mamelak, and Schuman (2006) provided evidence for novelty-detection neurons in the hippocampus by recording directly from hippocampal neurons using microwires in epilepsy patients. They were able to distinguish two different classes of neurons in the hippocampal-amygdaloid complex that exhibit single-trial changes in firing rates to novel versus familiar stimuli. Novelty-detectors showed increased firing rate in response to novel stimuli, while familiarity-detectors show increased firing in response to the second presentation of a previously encountered stimulus. The neuronal responses during recall did not differ between the first and second recognition phase (30 minutes versus 24 hours after the first stimulus presentation). This study shows that hippocampal neurons can reliably distinguish between novel and familiar stimuli for a prolonged amount of time.

Converging evidence from single unit recordings and ERP studies thus suggests that signals from the hippocampus can distinguish new from old. It remains unclear, however, whether a novelty signal also originates from the hippocampus, or whether hippocampal firing simply reflects novelty detection at an earlier processing stage. As it is known that the hippocampus does not actually store all previously encountered stimuli, it is possible that the hippocampus receives novelty-related inputs from other regions (Squire et al., 2004; Wiltgen, Brown, Talton, & Silva, 2004; Zhang & Zhu, 2011; Zola-Morgan & Squire, 1993). Probably because novelty detection and processing is crucial for survival and novelty is differently operationalized in different fields, ranging from extremely simple deviant sounds to complex environments (Schomaker, 2019), a large network of brain regions has been identified that responds to novelty. For example, novelty-related signals in the hippocampus could alternatively arise in the anterior cingulate cortex (i.e., signal reflecting prediction-errors or surprise) Alexander & Brown, 2019), ventral tegmental area (Lisman & Grace, 2005), or orbitofrontal cortex (Petrides, 2007). Within the MTL, also the perirhinal cortex has been associated with processing of stimulus novelty (Nyberg, 2005; Kumaran & Maguire, 2007), and the recognition of novelty (Scofield, Trantham-Davidson, Schwendt, Leong, Peters, See & Reicher, 2015). Another MTL structure, the amygdala, responds to novelty, and its response to stimuli habituates with repeated presentation (Blackford, Buckholtz, Avery, & Zald, 2010); Bradley, Lang, & Cuthbert, 1993; Kiehl et al., 2005; Schwartz et al., 2003).

In one previous study investigating the origin of a novelty signal, patients with posterior hippocampal damage were tested in an ERP design using and oddball paradigm with auditory and somatosensory stimuli (Knight, 1996). Interestingly, novelty detection, as indexed by the anterior N2, was unaffected by the MTL lesion, while the further processing of the novel stimuli, as indexed by the novelty P3, was reduced for low frequency stimuli labeled novel, presented contralateral to the resected side. These stimuli were not truly novel – they were what can be referred to as contextually novel and/or deviant (Schomaker & Meeter, 2015; Schomaker et al., 2014). The stimuli acting as novels could have previously been experienced by patients, including finger taps, electric shocks, and environmental sounds (like a bell), but were new within the context of the experiment. Moreover, in the Knight (1996) study the extent of the lesions differed substantially between patients, as the lesions were caused by an infarction of the posterior cerebral artery. Damage comprising the ‘hippocampus proper, dentate gyrus, subiculum, parahippocampal gyri, enthorhinal cortex and fornix in all patients’, but extended to the ‘fusiform, lingual and calcarine gyri’ in some patients and only less than half of the average lesioned regions showed overlap (Knight, 1996).

In the present study, a group of patients who underwent a unilateral MTL resection to treat their medically refractory temporal lobe epilepsy, was tested. The extent of the resections ranged from amygdalo-hippocampectomy to complete unilateral MTL resection. It is impossible to create simple auditory or somatosensory stimuli that are truly novel (Schomaker & Meeter 2015). For this reason, in contrast with the study by Knight (1996), the patients performed a visual rather than auditory or somatic version of the novelty oddball task while their electroencephalogram was measured. Moreover, truly novel rather than deviant stimuli were employed, consisting of line-drawings of novel objects that the participants never saw before the experiment, nor were they repeated within the experiment. The visual novelty oddball task was adapted such that novel and standard stimuli were either presented contra- or ipsilateral to the subjects’ lesion, while a visually matched standard stimulus was presented on the opposite side. This design using lateralized stimuli allowed us to investigate the role of the MTL in the detection and processing of novelty in a within-subjects design. Stimuli that are presented on one side are mainly processed by the contralateral hemisphere. This means that a stimulus presented ipsilateral to the resected side, is processed by the intact hemisphere, while a stimulus contralateral to the resection would be processed by the affected hemisphere. Note, that the visual cortex was intact on both sides, therefore the current study allowed us to investigate the role of the MTL in visual novelty processing. Exploiting the high temporal resolution of the ERP technique we aimed to identify the role of MTL structures in the detection and later evaluation of visual novelty. We expected that novelty detection, as measured by the novelty N2, and/or further processing of novelty, as measured by the P3, would be reduced for novel stimuli presented to the resected compared to the unresected side. We did not expect target processing to be altered.

## Methods

### Participants

Twenty-one patients who underwent a MTL resection to treat drug-resistant focal epilepsy volunteered to participate (8 female; age range: 19-64; mean age 42.5 years; standard deviation [SD] age = 14.6 years; 19 right-handed). Nine patients had undergone a left temporal lobe resection (2 female; age range: 20-60; mean age 38.7 years; SD age = 13.8 years; 8 right-handed), and twelve patients had undergone a right temporal lobe resection (6 female; age range: 19-64; mean age 44.8 years; standard deviation [SD] age = 14.5 years; 11 right-handed). The patients were on average 578 days (range 345-931 days) post-surgery. All participants received an informational invitation letter and signed informed consent before starting the experiment. All patients had undergone unilateral resection of the anterior temporal lobe including the hippocampus in the hemisphere of the epileptic focus. A standard temporal resection extends on the lateral side up to 3 to 4 cm, while a maximal temporal resection on the non-language dominant side extends up to 8 cm from the temporal pole. On the language dominant side, the extend of a maximal resection was limited by the location of Wernicke’s area, as determined either with an awake procedure or Penfield procedure or during testing with a grid-implantation. All patients were taking anti-epileptic medication at the time of participation and they all were native Dutch speakers. Patient descriptions are summarized in Table 1.

**Table 1.**
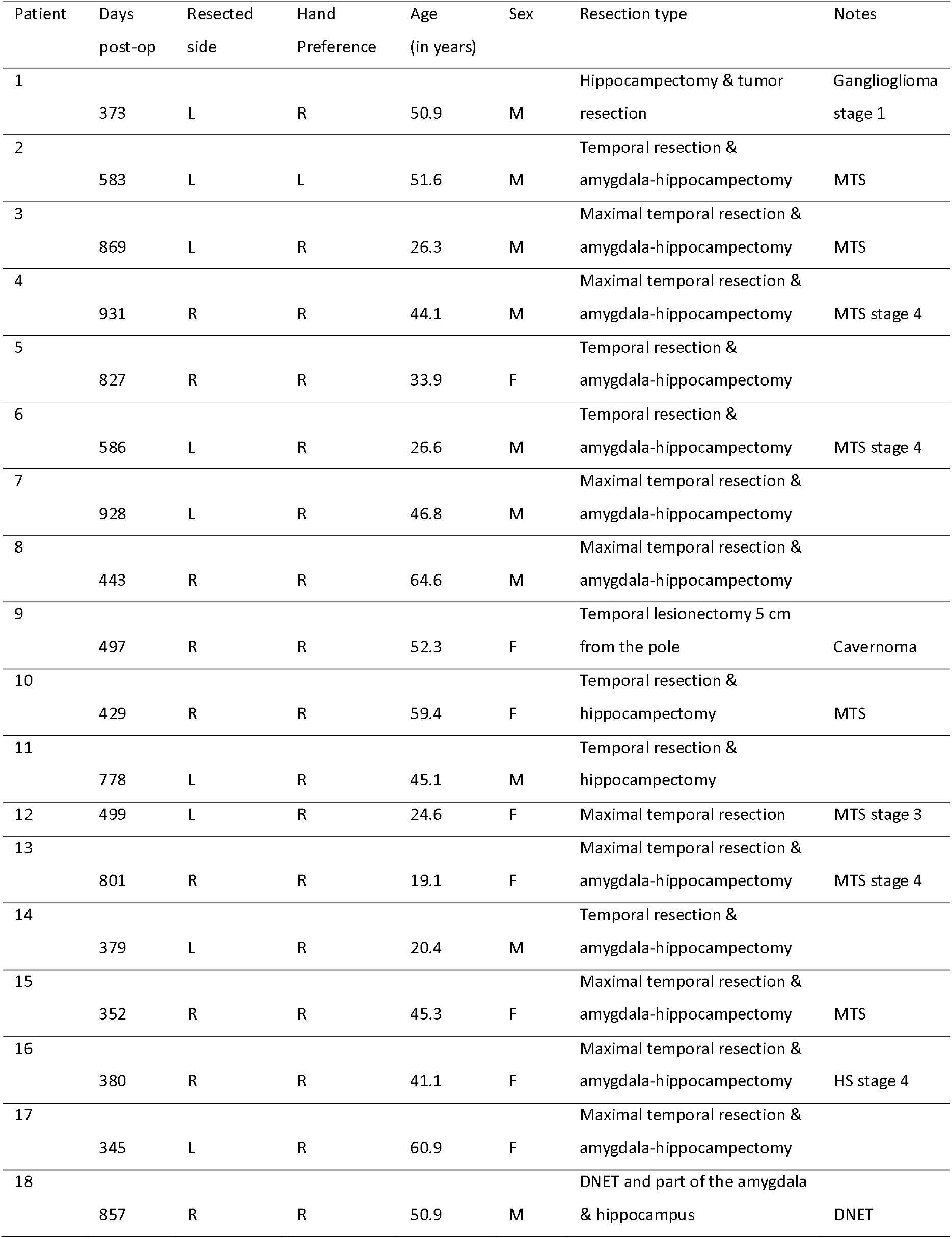
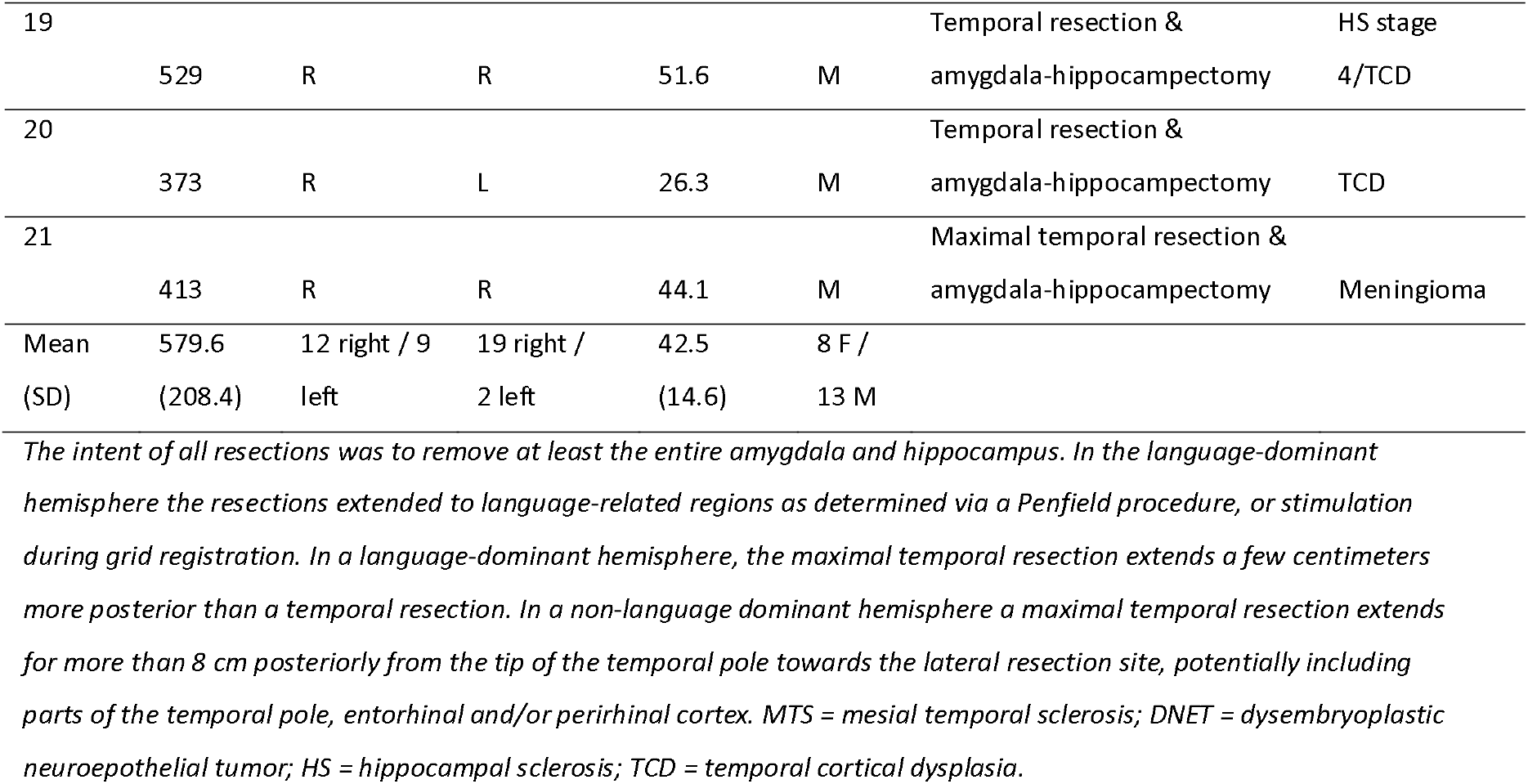
Patient details

After study completion participants were debriefed about the major study’s findings. This study was approved by the Committee of Research and Development and the Medical Ethical Permission Committee of the Epilepsy Centre Kempenhaeghe, the Netherlands. In accordance with the ethical guidelines the study was performed during a routine check-up EEG measurement, and participants received no financial compensation.

### Stimuli

There were three types of stimuli, novels, standards, and target oddballs. Novel stimuli consisted of white line-drawings of non-existent objects (gracefully provided by Kirk Daffner, and also used in for example Chong et al., 2008), and were not repeated throughout the experiment. Standards were scatter plots created by randomly moving around the white pixels of the novel stimuli within the boundaries of a circle. This was done to create standard stimuli matched to the novels and target in terms of luminance (i.e. scrambling the white pixels of novels and presenting them in a circular shape on a black background; also see Schomaker et al., 2014). When a novel or target stimulus was presented its matching circular scattered version was presented on the opposite side. Targets were triangles that could point upwards or downwards (counterbalanced between subjects). All stimuli were covered about 3.8}3.0 cm and were presented at a distance of 12.5 cm on a black background.

### Experimental Procedure

Patients were seated in a dark EEG recording room, and participants performed the novelty oddball task while their EEG was measured. A pilot study in non-matched students (findings not reported here) confirmed that this task induced the typical novelty N2 and P3 components. Stimuli were presented on a 15.4 inch laptop with a 1440×900 pixel screen using E-Prime version 2.0 (Psychology Software Tools). Viewing distance was about 60 cm. They were instructed to minimize blinking and motion, and fixate the fixation cross whenever it was presented. All subjects were given a practice block consisting of 20 trials. On each trial, two visual stimuli were shown together; one stimulus was presented on the left side of a fixation cross and the other on the right side. On standard trials the same standard stimulus was shown on the left and right of fixation. Novels and targets were shown either on the left or right during a specific trial, while a standard, created by shuffling the pixels of the simultaneously presented novel/target, was shown on the opposite side. See Figure 1 for example stimuli. Standard trials were frequent (71.4%), while novel and target trials were infrequent (14.3%). Participants were instructed to press the letter ‘b’ with the right or left index finger (counterbalanced across subjects) as quickly as possible, irrespective of whether the target stimulus appeared on the left or right side of a fixation cross. On each trial the stimuli were presented for 2000 ms. Between trials a fixation cross was shown for 800-1500 ms.

**Figure 1.**
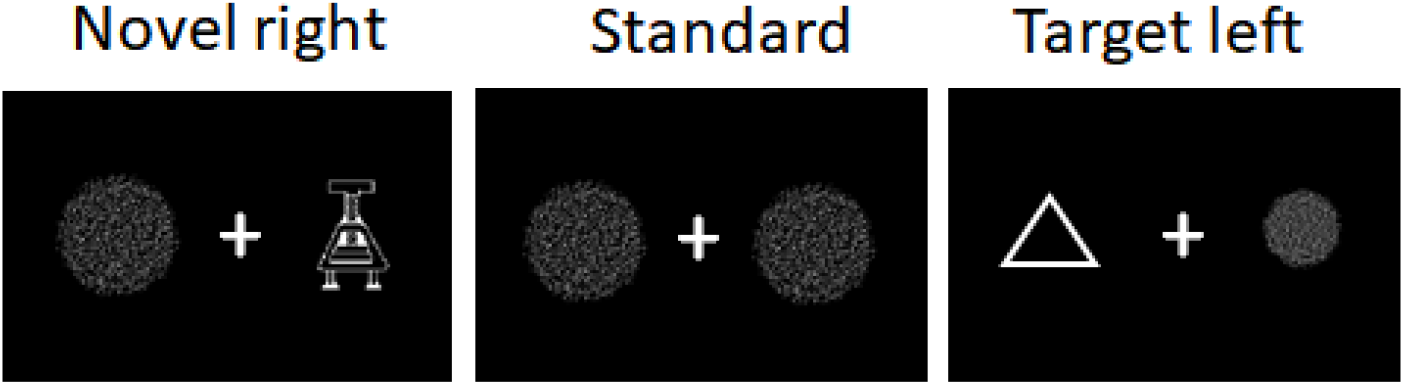
Example stimulus combinations. On the left a display is shown with a novel stimulus presented on the right side with its scattered circular counterpart on the left. In the middle an example standard display is shown, with two matched scatter circles. On the right a display is shown with a target on the left and its scattered circular counterpart on the right. Participants merely had to detect the target and respond with a button press. Stimulus displays were shown for 2000 ms, and between trials a fixation cross was shown for a jittered interval of 800 to 1500 ms. Note that pictures were rescaled for demonstrational purposes in this figure.

The experimental task was divided into six blocks that consisted of 70 stimuli each. Each block had a duration of approximately 5 minutes. Between blocks participants could take a self-paced break. The experimental procedure had a total duration of about 35 minutes.

### EEG recording

All silver disc Ag Grass electrodes were placed (Grass Technologies) on the scalp. The electrodes used to measure horizontal (F9 and F10) and vertical (SO1 and SO2) electrooculogram (EOG) were placed respectively at the outer canthi of both eyes and supraorbitally. EEG Ag/AgCl electrodes were placed on the scalp with an Electrocap (Electrocap International, Inc) using the 10/20 system developed by the International Federation of Clinical Neurophysiology (Jasper, 1958). The electrodes placed on the scalp were at 10 and 20 percent of the lines linking these landmarks. Impedances was kept below 10kΩ. Data was recorded using BrainRT version 1.0.1972 with a bandpass of 0.3 to 70 Hz, and digitized at 250 Hz for offline analysis. An EDF converter (www.OSG.be) was used to transform BrainRT files into EDF format. Since data was collected in a clinical setting, additional reference electrodes were not used in all participants, therefore, recordings were offline rereferenced to the average of all 22 EEG electrodes (Fp1, Fpz, Fp2, F9, F7, F3, Fz, F4, F6, F8, T3, C3, Cz, C4, T4, T5, P3, Pz, P4, T6, O1, O2). Data was offline re-referenced to the average of all. Data analyses were performed on the remainder of the data.

### EEG analyses

Data was analysed using EEGlab toolbox version 13_5_4b (Delorme & Makeig, 2004) and the BioSig Toolbox (OSG). Epochs were created for a time window of −200 to 1500 ms around stimulus onset. The data was visually inspected by a trained expert to identify and remove trials with motion artefacts, eye movements and blinks within the N2-P3 time-window in the electrodes of interest (Fz, Cz, and Pz). As we used these strict criteria, we were able to conserve a relatively large part of the data. For novels and targets a minimum of 15 and a maximum of 30 epochs (mean = 29.2 epochs, with a maximum of 30) were included per condition (ipsi- and contralateral) per participant. For standards a minimum of 257 and a maximum of 300 was included. ERPs were calculated for the remainder of the data relative to a −200 to 0 ms prestimulus baseline using standard signal averaging techniques (Luck, 2005). Mean amplitudes were calculated for the components of interest for each of the conditions at three different electrode sites (Fz, Cz, and Pz). Based upon visual inspection of the grand-average ERP signal, mean amplitudes were calculated for a time-window of 190-240 ms for the N2, 290-340 ms for the P3, and 275-450 for the target P3b.

### Statistical analysis

Behavioral results were investigated with a repeated-measures analysis of variance (ANOVA) with Target location (left; right) as within-subjects factor and Resected side (left; right) as a between-subjects factor.

The N2 and novelty P3 elicited by novels were investigated with a repeated-measures ANOVA with Resected side (ipsilateral; contralateral) and Electrode (Fz; Cz; Pz) as within-subjects factors. Ipsilateral defined as a stimulus presented on the same side as the resection, and contralateral as a stimulus presented opposite to the resected side.

In two separate ANOVAs ERPs elicited by the different stimuli were compared, to confirm that our lateralized version of the visual oddball task elicited typical novelty ERP responses. Differences between novels and targets were investigated with a repeated-measures ANOVA with Stimulus (novel; target), Resected side (ipsilateral; contralateral) and Electrode (Fz; Cz; Pz) as within-subjects factors. Differences between novels and standards were investigated with a repeated-measures ANOVA with Stimulus (novel; standard), and Electrode (Fz; Cz; Pz) as within-subjects factors. Note that standards were always presented on both the left and right side, therefore we collapsed over ipsi- and contralateral conditions for the novels for this analysis. For this ANOVA we only report the Stimulus main effect, interactions with Stimulus, and corresponding post-hoc comparisons.

Since standards elicited a smaller N2 than novels, this also affected the size of the following positive-going component, by keeping it on the positive side of the scale. To control for these early stimulus differences, we also performed an ANOVA with Resected side (ipsilateral; contralateral) and Electrode (Fz; Cz; Pz) as within-subjects factor for the novel-elicited minus the standard ERP components. For all ANOVAs Greenhouse-Geisser correction was performed when the sphericity assumption was violated.

## Results

### Behavioral responses

Participants with a left MTL resection responded to the visual target accurately on 97.88% (standard deviation [SD] = 7.28%), and participants with a right MTL resection on 98.41% (SD = 5.46%) of target-present trials. Hit rate did not differ between the two groups (p = .300), nor was there an interaction effect between resected side and target location (p = .390). Very few false alarms were given, and the false alarm rate did not differ between the two patient groups (p = .869), with 0.69% (SD = 1.36) false alarms for patients with left and 0.76% for patients with right MTL resections. Figure 2 shows box plots and individual mean response times for patients with left- and right-sided MTL resections for ipsi- and contralaterally presented targets. Response times also did not differ between the two patient groups, with a mean response time of 544 ms (SD= 141 ms) for patients with left and 576 ms (SD = 131 ms) for patients with right MTL resections (p = .302). However, resected side interacted with target location, F(1,19) = 8.14, p = .010, η = .30, with patients with a right MTL resection responding faster for targets presented on the right rather than left (57 ms difference), and the reverse for patients with a left MTL resection (25 ms difference). This suggests that patients were faster in detecting and responding to a target presented on the ipsilateral side, i.e., to their unaffected hemisphere.

**Figure 2.**
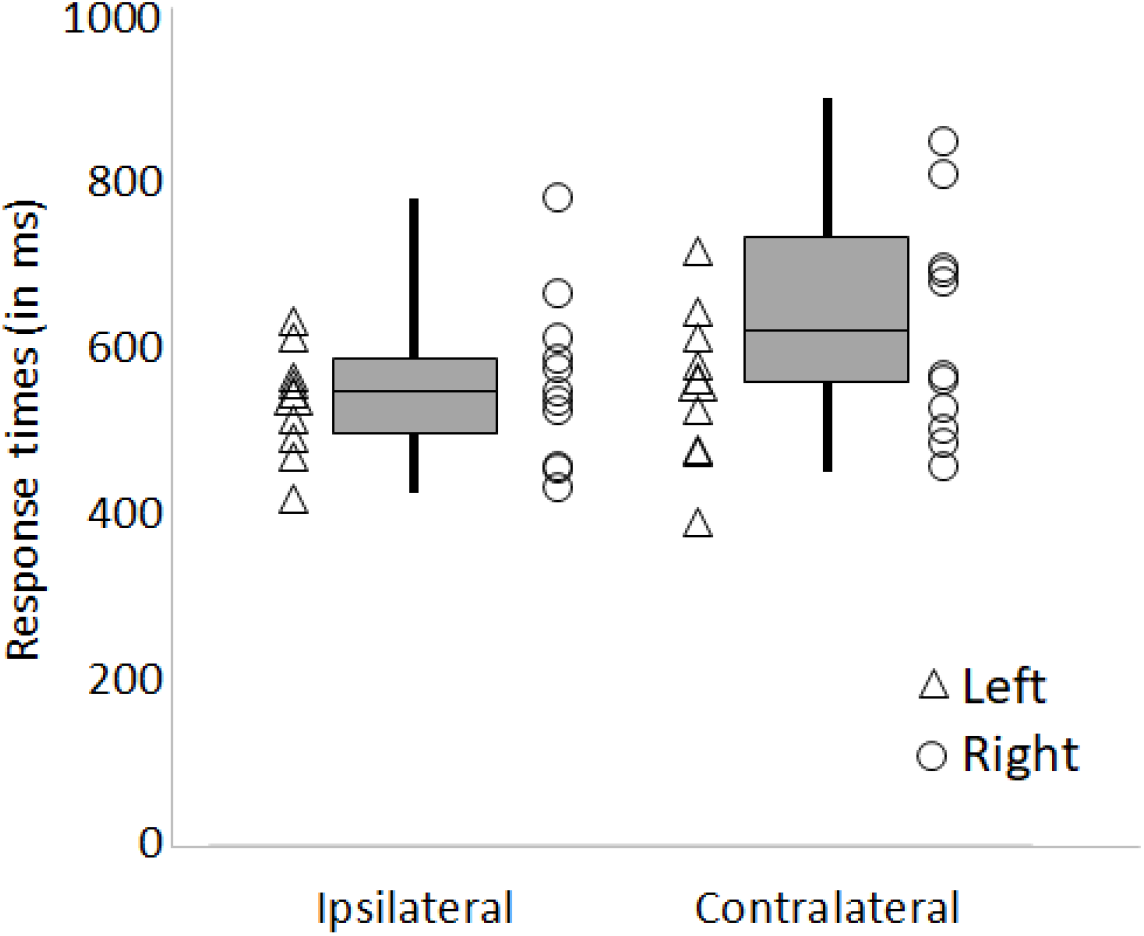
Box plots and individual mean response times in milliseconds for patients with left-(triangles) and right-sided (circles) MTL resections for ipsi- and contralaterally presented targets.

### Mean amplitude ERPs

Novels elicited a frontal negative component (N2) peaking around 220 ms, followed by a positive going wave(P3) peaking slightly later over posterior electrodes. Figure 3 shows the grand-average ERPs for ipsi- and contralaterally presented novels, and bilaterally presented standards, and ipsi- and contralaterally presented targets. Figure 4 shows mean N2 and P3 peaks for individual patients for ipsi- and contralterally presented novels. Figure 5 shows the topographic plots for ipsi- and contralaterally presented novels and bilaterally presented standards in the N2 and P3 time-windows. Figure 6 shows the topographic maps for ipsi- and contralaterally presented targets in the P3b time-window.

**Figure 3.**
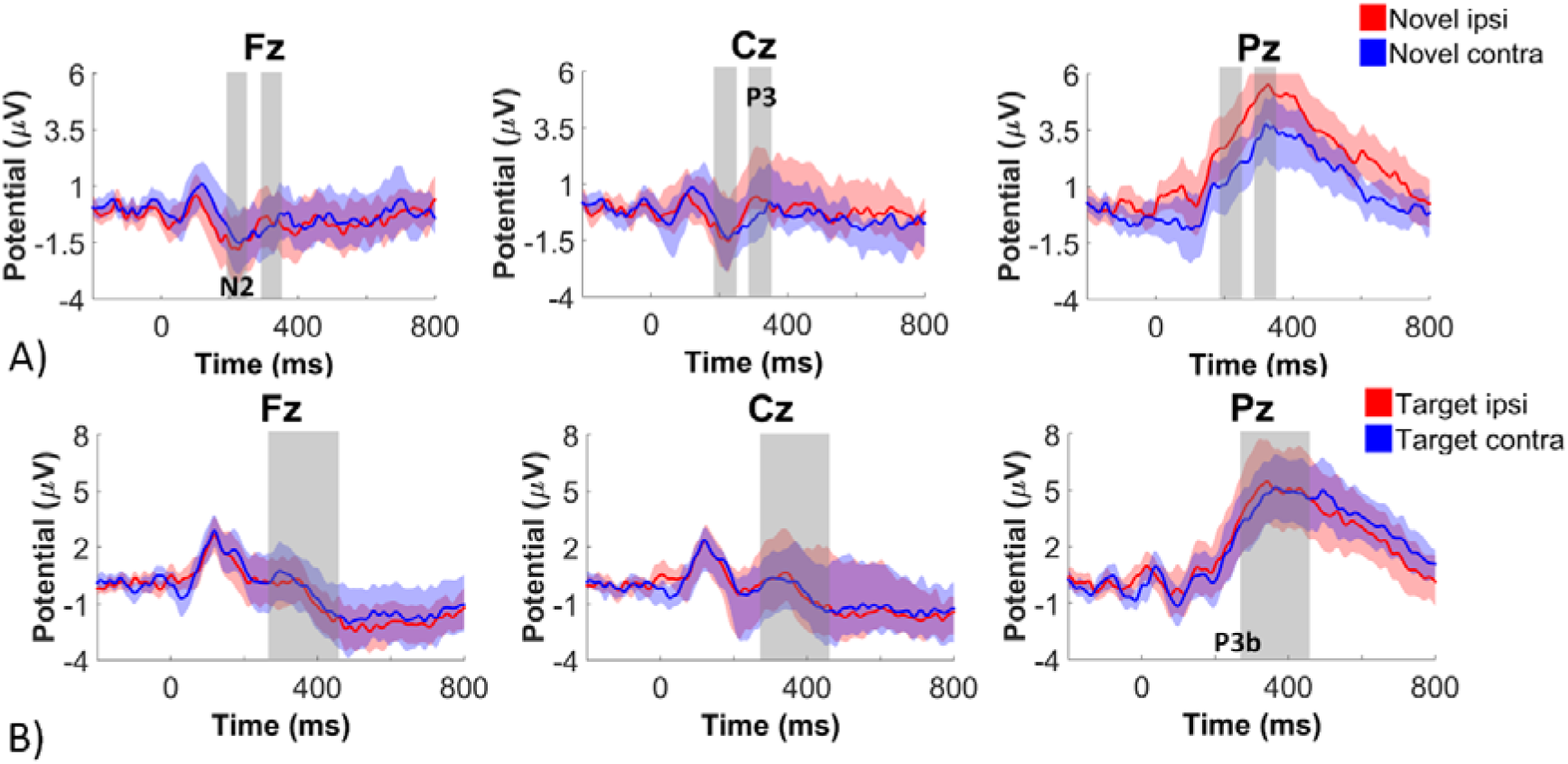
A) ERPs for novels presented ipsi- and contralaterally at Fz, Cz and Pz, and B) ipsi- and contralaterally presented targets atFz, Cz, and Pz. The time-windows used for the mean amplitude ERP analyses are highlighted in grey for the N2 and P3 components (A), and the P3b component (B). The translucent areas depict 95% intervals around the means per stimulus condition.

**Figure 4.**
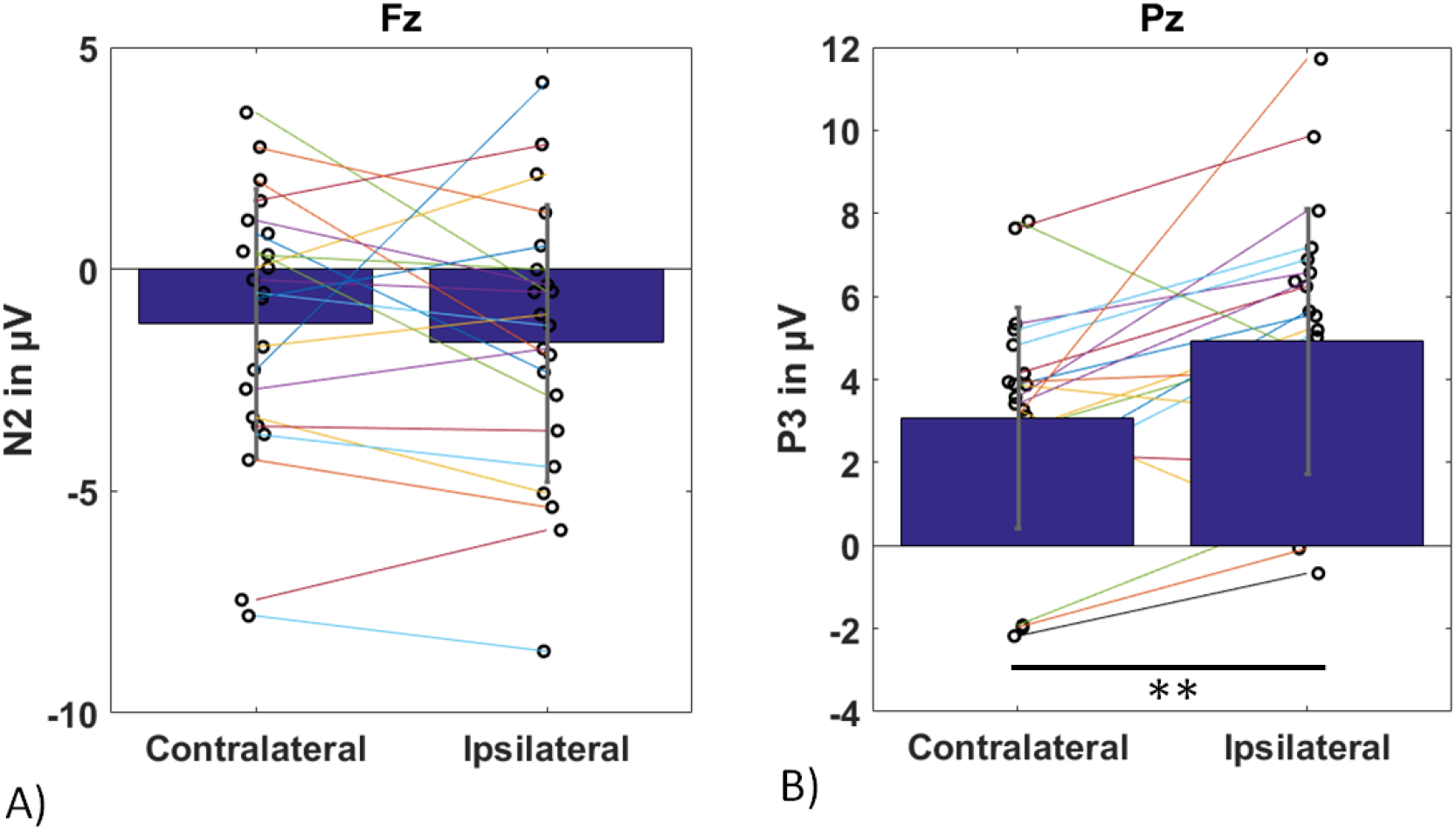
Mean and individual mean ERP amplitudes for A) the N2 component at Fz, and B) the P3 at Pz for contralateral and ipsilateral novels.

**Figure 5.**
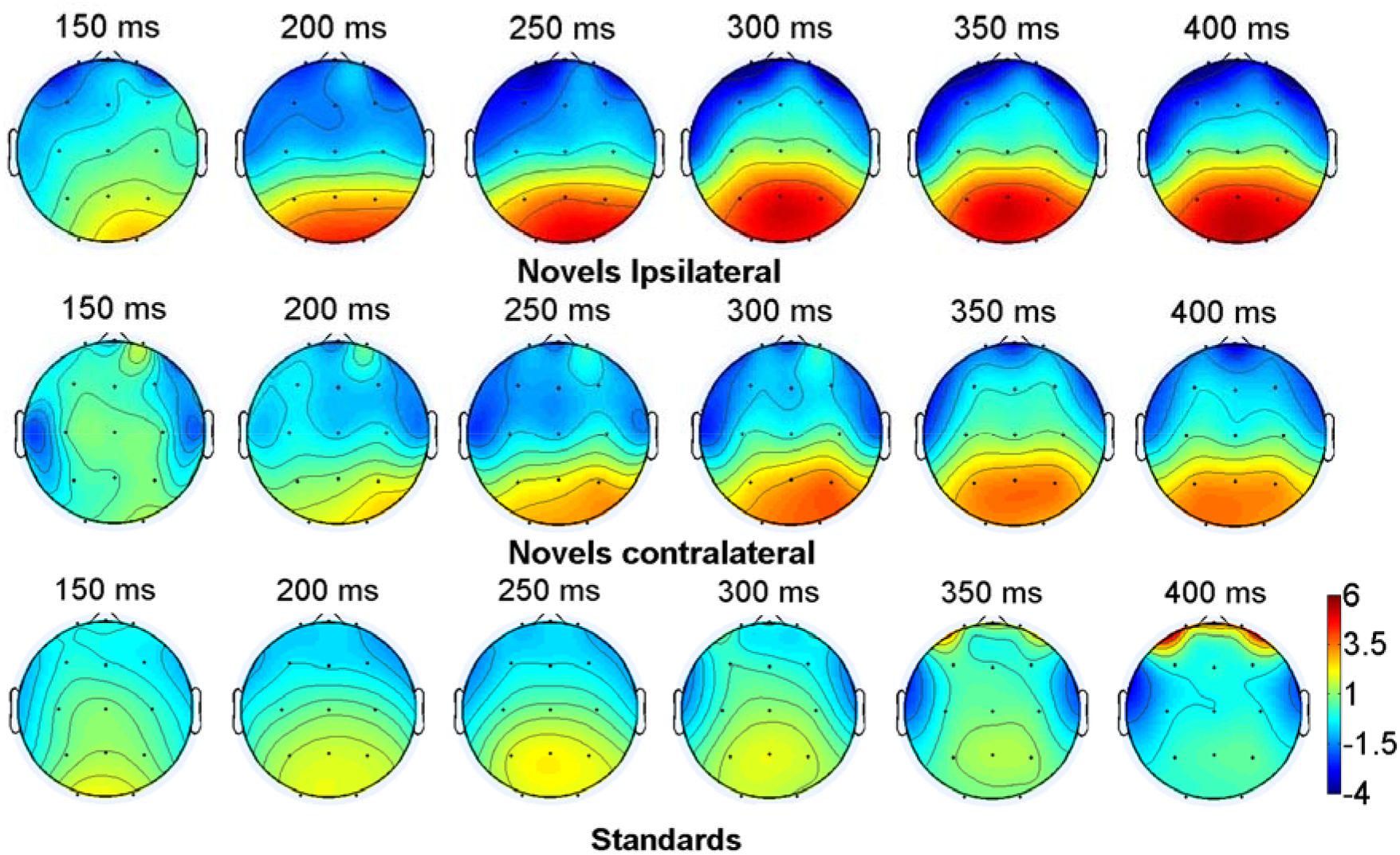
Topographic maps for novels presented ipsilaterally (top) and contralaterally (middle) and standards (bottom). Topographic maps are shown for 150, 200, 250, 300, 350, and 400 milliseconds post-stimulus. In these time-windows the N2 and P3 components peaked.

**Figure 6.**
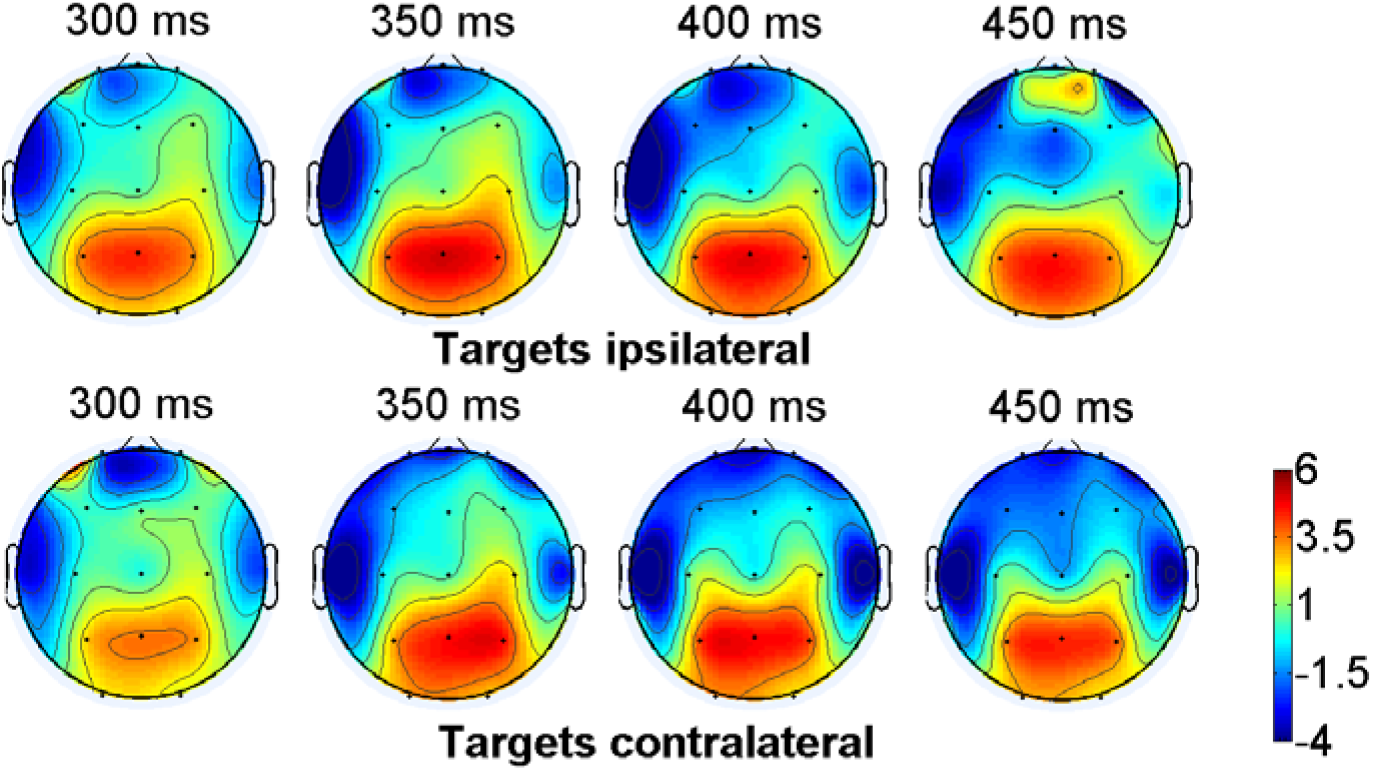
Topographic maps for targets presented ipsilaterally (top) and contralaterally (bottom). Maps are shown for 300, 350, 400, and 450 milliseconds post-stimulus. In this time-window the P3b component peaked.

### Anterior N2

The anterior N2 was investigated with a repeated-measures ANOVA with Resected side (ipsilateral; contralateral) and Electrode (Fz; Cz; Pz) as within-subjects factors. For novel stimuli there was no main effect of Resected side on N2 amplitude (p = .320; also see Figures 3 and 4). The N2 peaked over frontal regions as shown by a linear effect of Electrode, F(1,20) = 15.56, p = .001, η = .44. Resected side and Electrode interacted, F(1.46,29.27) = 5.99, p = .012, η =.23. We further investigated this interaction with a t-test comparing the anterior N2 at Fz for contralateral and ipsilateral presented novels, but no difference was observed (p = .429).

A separate ANOVA investigating differences between standard- and novel-evoked ERPs was performed. A statistical trend suggested that novels elicited a larger N2 than standards, F(1,20) = 3.93, p = .061, η = .16. Stimulus also interacted with Electrode, F(2,40) = 9.20, p = .001. Post-hoc t-tests to identify the peak location, compared the N2 for novels and standards. The N2 was larger for novels than standards at Cz, t(20) = 2.84, p = .010. For Fz there was trend effect, t(20) = 2.09, p = .050, but this did not survive Bonferroni correction. Posteriorly no differences in standard- and novel-evoked N2 were found (p = .716).

### Novelty P3

Figure 4B shows mean P3 peaks at Pz for ipsi- and contralaterally presented novels for individual patients. As for the anterior N2 the effects of resected side and electrode were investigated for the novelty P3. Novel stimuli elicited a larger novelty P3 when presented ipsilateral compared to contralateral to the resection, F(1,20) = 5.24, p = .033, η = .21. The novelty P3 peaked over posterior regions as shown by a linear effect of Electrode, F(1,20) = 27.60, p < .001, η = .58.

Resection side and Electrode also interacted, F(1.54,30.74) = 5.15, p = .010, η =.21. Post-hoc tests showed that the novelty P3 was larger for ipsilateral than contralateral presented novels at Pz, t(20) = 3.38, p = .003. No differences were observed on the other electrodes (ps > .101).

Also the stimulus effect was investigated again. Novels elicited a larger P3 than standards, F(1,20) = 5.32, p = .032, η = .21. Stimulus also interacted with Electrode, F(1.19,23.75) = 11.73, p < .001. Post-hoc t-tests comparing the P3 for novels and standards showed that the P3 for novels was larger at Pz, t(20) = 4.19, p < .001, while no differences were observed at Fz (p = .117) or Cz (p = .983).

To further isolate the effects specific to novelty, the same analyses were performed on the difference waves for the novelty P3 component elicited by novel minus standard,. In line with the main novelty P3 analyses, novel stimuli elicited a larger novelty P3 when presented ipsilateral compared to contralateral to the resection side when subtracting the standard-elicited P3, F(1,20) = 5.24, p = .033, η = .21. The novelty P3 peaked over posterior regions as shown by a linear effect of Electrode, F(1,20) = 12.35, p = .002, η = .38. Resection side and Electrode also interacted, F(1.54,30.74) = 5.15, p = .010, η =.21. Post-hoc tests showed that the novelty P3 was larger for ipsilateral than contralateral presented novels at Pz, t(20) = 3.38, p = .003 (also see Figures 3 and 4). No differences were observed on the other electrodes (ps > .101).

### Target P3b

To check whether the MTL resections affected target processing, we investigated the P3b with a repeated-measures ANOVA with Resected side (ipsilateral; contralateral) and Electrode (Fz; Cz; Pz) as within-subjects factors. The P3b was of similar size when targets (i.e. oddballs) were presented ipsi- and contralateral to the resected side (p = .703). The P3b peaked over posterior regions as shown by a linear effect of Electrode, F(1,20) = 16.26, p = .001, η = .45 (see Figure 6). Resected side and Electrode did not interact (p = .314).

## Discussion

Epilepsy patients with unilateral MTL resections performed an adapted version of the visual novelty oddball task. Novel and target stimuli were either presented to the resected (contralateral) or to the unresected (ipsilateral) side. Target processing was not affected by this manipulation. Early processing of novelty was not affected by the MTL resections either, but the later processing was reduced for novels presented to the resected compared to the intact side.

The MTL has been suggested to be the origin of a novelty signal (Knight, 1996; Kumaran & Maguire, 2007; Ranganath & Rainer, 2003). The N2 elicited by novels is believed to be related to the perceptual part of novelty detection, and is believed to reflect an early novelty signal (Ferrari et al., 2010; Schomaker et al., 2014). In the main analyses no differences in anterior N2 amplitude were observed for ipsi- and contralaterally presented novels, suggesting that novelty detection is not affected by the MTL resection: At its frontal peak location, no differences were observed. These findings are in line with a previous study, suggesting that novelty detection does not take place in the MTL (Knight, 1996). Another possible site for novelty detection is the lingual gyrus, which lies in the medial ventral visual stream, posteriorly to the MTL. Using neuroimaging it has been linked to perceptual processing of unattended novel stimuli (Stoppel et al., 2009). It has previously been suggested as a possible source of the novelty N2 component, which was proposed to reflect stronger neural firing to stimuli that do not yet have a sharpened perceptual representation (Schomaker & Meeter, 2015). Also the superior colliculus has been suggested to carry an early novelty signal (Aito et al., 2007; Boehnke et al., 2011).

In contrast with the N2, the later novelty P3 component was reduced for novels presented contra-versus ipsilateral to the resected side. This suggests that a novelty or deviance signal is coming from the MTL. Importantly, the current findings cannot be explained by a general stimulus processing detriment, as the processing of ipsi- and contralaterally presented target information was not differently processed as evidenced by a similar size P3b. It has been proposed that the novelty signal generated by the hippocampus is the result of violated predictions (Kumaran & Maguire, 2007, 2009). The ‘novelty signal’ in the MTL may thus reflect detection of stimuli that are unexpected or deviant stimuli in the current context, and perhaps should be referred to as a deviance signal instead. For example, using fMRI Strange and Dolan (2001) found that the anterior hippocampus responds to perceptual, semantic, and emotional oddballs, and argued that it detects a general mismatch between expectations and outcome, rather than novelty per se. Furthermore, hippocampal and surrounding MTL activity is reduced with continued exposure to novelty (Murty et al., 2013). These studies suggest that the hippocampus does not simply code for novelty, but dynamically responds to the presence of novelty in the environment by adjusting the allocation of neural resources accordingly. Interestingly, also the novelty P3 has been linked to the frequency and deviance of novel stimuli, and the resulting effects on expectations (Cycowicz & Friedman, 2007; Schomaker, Meeter (2018); Schomaker et al., 2014). The current study’s results are in line with the idea that the hippocampus responds to mismatch with expectations, rather than novelty per se, however, future studies are required to address this question more directly.

The finding that the P3b was not affected by the MTL resections is in line with the findings of Knight (1996), but in contrast with other studies that suggested that the hippocampus is the main generator of the P3b component (Halgren, Marinkovic, & Chauvel, 1998; McCarthy, Wood, Williamson & Spencer, 1989; Halgren, Squires, Wilson, Rohrbaugh, Babb & Crandall, 1980). This discrepancy may have been caused by differences in the oddball tasks used. Knight (1996) and the present study employed oddball tasks with a third category of stimuli (novels/deviants), while studies that identified hippocampal involvement in the P3b, mainly used two-stimulus oddball tasks to study the P3b component, however, future work is needed to further investigate this possibility. Although we did not observe differences in P3b amplitudes, participants were actually slower in detecting the target when it was presented to the resected (contralateral) versus unresected (ipsilateral) side in the present study. Numerically, there was a slight slowing of P3b peak for the contra-relative to ipsilaterally presented targets, so MTL resections might have slowed down – though not altered – target processing, or hampered response preparation processes in the current study.

Novel stimuli typically activate a widespread network in the brain, including frontal, temporal visual, and subcortical regions (Blackford, Buckholtz, Avery, & Zald, 2010; Bunzeck & Düzel, 2006; Daffner et al., 2000; Ranganath & Rainer, 2003; Tulving, Markowitsch, Kapur, Habib, & Houle, 1994; Wendt, Weike, Lotze, & Hamm, 2011; Wright et al., 2003). Similarly, various regions have been associated with the P300 component (Knight, 1984; Ludowig, Bien, Elger, & Rosburg, 2010). It is therefore likely that a network of regions, rather than a single region underlie novelty detection and processing. The results of the current study do suggest however, that MTL structures including the hippocampus and amygdala play some role in the complex process of identifying, processing, and storing novel information. One limitation of the current study is the lack of a healthy control group. Note, that we used a lateralized task in a within-subjects design that allowed us to make comparisons between information processed in the resected and non-resected hemispheres. As would be expected, we observed the typical N2 and P3 novelty components in the to-be-expected time-windows, suggesting that our lateralized task was effective in eliciting typical novelty ERP components. Most interestingly, we observed differences between ipsi- and contralateral presented stimuli: Reduced novelty processing in the hemisphere with MTL resection. The lack of a control group, however, limits the generalizability of our findings.

In the patients of the current study, MTL regions were surgically removed using anatomical landmarks. Each surgery was aimed at completely removing the hippocampus and amygdala, resulting in very similar resection patterns. Although the patient sample was more homogeneous as in the Knight (1996) study, in which lesions occurred naturally and sometimes extended to other regions, another limitation of the current study is that the patients’ MTL resections included not only the hippocampus, but also the adjacent amygdala in 17 out of 21 patients. Also the extent of the resections varied between patients (see Table 1). In the case of a maximal temporal resection parts of the temporal pole, entorhinal cortex and perirhinal cortex were removed as well. Variation was insufficient, however, to divide the patients into groups or investigate individual differences as all patients had overlap between the resected regions.

The reduced novelty P3 for novels presented contra-versus ipsilateral to the resection could be caused by the absence of a hippocampus on the contralateral side, however, novelty also activates other MTL structures. For example, the amygdala has been known to code for novelty, and the amygdala response to emotional stimuli habituates with repeated presentation (Blackford et al., 2010; Bradley, Lang, & Cuthbert, 1993; Kiehl et al., 2005; Schwartz et al., 2003), possibly signaling the salience of new stimuli. Even when separating effects of arousal, valence and novelty, novelty has been found to be a relevant driver of activity in the affective circuits in the human brain (Weierich, Wright, Negreira, Dickerson, & Barrett, 2010). The current study’s results could therefore be explained by the patients’ amygdala resections, or the strong connection between the hippocampus and amygdala (Fastenrath et al., 2014). Previous work also suggested that the perirhinal cortex plays a role in novelty processing (e.g., Nyberg, 2005; Kumaran & Maguire, 2007; Scofield, et al., 2015). Especially in patients with maximal temporal resections parts of the perirhinal cortex may have been removed. Removal of part of the perirhinal cortex in the 10 patients that underwent maximal temporal resection therefore may partly underlie the current study’s results.

In any case, our findings confirm that MTL structures, including the hippocampus and amygdala, may play a role in novelty processing, but are not required for the earlier, perceptual processing or detection of novelty.

